# Deep Residual Learning for Neuroimaging: An application to Predict Progression to Alzheimer’s Disease

**DOI:** 10.1101/470252

**Authors:** Anees Abrol, Manish Bhattarai, Alex Fedorov, Yuhui Du, Sergey Plis, Vince D. Calhoun, for the Alzheimer’s Disease Neuroimaging Initiative

## Abstract

This work investigates the suitability of deep residual neural networks (ResNets) for studying neuroimaging data in the specific application of predicting progression from mild cognitive impairment (MCI) to Alzheimer’s disease (AD). We focus on predicting the subset of MCI individuals that would progress to AD within three years (progressive MCI) and the subset of MCI individuals that do not progress to AD within this period (stable MCI). This prediction was conducted first as a standard binary classification task by training a ResNet architecture using MCI individuals only, followed by a modified domain transfer learning version that additionally trained on the AD and cognitively normal (CN) individuals. For this modified inter-MCI classification task, the ResNet architecture achieved a significant performance improvement over the classical support vector machine and the stacked autoencoder machine learning frameworks (*p* < 0.005), numerically better than state-of-the-art performance in predicting progression to AD using structural MRI data alone (> 7% than the second-best performing method) and within 1% of the state-of-the-art performance considering learning using multiple structural modalities as well. The learnt predictive models in this modified classification task showed highly similar peak activations, significant correspondence of which in the medial temporal lobe and other areas could be established with previous reports in AD literature, thus further validating our findings. Our results highlight the possibility of early identification of modifiable risk factors for understanding progression to AD using similar advanced deep learning architectures.

## Introduction

Dementia is vastly underdiagnosed in most health systems mainly due to lack of educational/awareness programs and accessibility to dementia diagnostic, treatment and care services (Bradford et al. 2009; Connolly et al. 2011; Wilkins et al. 2007). Diagnosis typically occurs at relatively late stages, following which the prognosis is poor in most cases since even state of the art (FDA-approved) medications in these stages are, at best, only modestly effective in alleviating cognitive and behavioral symptoms of the disease. As such, early therapeutic interventions can not only help improve the cognitive and behavioral function of the elderly patients, but also empower them to take important decisions about their health care while they can, and significantly improve their overall quality of life.

The most widely reported form of dementia in the elderly population is Alzheimer’s disease (AD) that features progressive, irreversible deterioration in memory, cognition and behavioral function. Mild cognitive impairment (MCI) has been identified as an intermediate condition between typical age-related cognitive decline and dementia (Markesbery 2010). This condition often leads to some form of dementia (not necessarily AD) and hence is commonly referred to as the prodromal stage of the disease. However, in the absence of an exact (i.e. narrower) prodrome for AD, this broader population of MCI is currently an attractive target for testing preventive treatments of AD. As mentioned before, the presently approved preventive medications are effective only over a limited (early) period (Casey, Antimisiaris, and O’Brien 2010). As such, the modest effectiveness and extremely high costs of these drugs have been a matter of constant debate especially in terms of cost to benefit balance. Hence patients showing MCI symptoms must ideally be diagnosed at early stages and be followed up regularly to identify potential risks of progression to AD (or other types of dementia). Several studies are currently focused in this direction with a remarkable increase in the collection and processing of multimodal neuroimaging, genetic, and clinical data. As a straightforward example, there are as many as thirty-four different live datasets that can be accessed from the Global Alzheimer’s Association Interactive Network (GAAIN) funded by the Alzheimer’s Association (GAAIN Data 2017). Today, out of these splendid data collection efforts, it is primarily the longitudinal studies that act as a bridge between clinical and neuropathological models (Markesbery 2010).

The structural magnetic resonance imaging (sMRI) neuroimaging modality enables tracing of brain damage (atrophy, tumors and lesions) and assists in ruling out any possible causes of dementia other than AD. This modality has additional advantages for its non-invasive nature, high spatial resolution, and ease of availability. Over the last two decades, several studies have contributed to the identification of potential AD biomarkers and prediction of progression to AD using sMRI data independently or in a multimodal pipeline (Arbabshirani et al. 2017; Falahati, Westman, and Simmons 2014; Rathore et al. 2017; Weiner et al. 2017). At the same time, the neuroimaging community has increasingly witnessed successful application of standard (i.e. classical) and advanced (i.e. deep or hierarchical) machine learning (ML) approaches to extract discriminative and diagnostic information from the high dimensional neuroimaging data (Litjens et al. 2017; Plis et al. 2014; Shen, Wu, and Suk 2017; Vieira, Pinaya, and Mechelli 2017). ML approaches are being increasingly preferred also because they allow for information extraction at the level of the individual thus making them capable of assisting the investigator in diagnostic and prognostic decision-making of the patients. The ML methods could range from standard classification frameworks (for example, logistic regression or support vector machines) that usually require manual feature engineering as a preliminary step to deep learning architectures that automatically learn optimal data representations through a series of non-linear transformations on the input data space. The last few years have seen an emergence of deep structured or hierarchical computational learning architectures to learn data representations that enable classification of brain disorders as well as predicting cognitive decline. These architectures hierarchically learn multiple levels of abstract data representations at the multiple cascaded layers, making them more suitable to determine subtle differences in the data. Some popular deep learning architectures including multilayer perceptron, autoencoders, deep belief nets, and convolutional neural networks have indeed been applied for AD classification and predicting progression of MCI patients to AD (Chen et al. 2015; Falahati, Westman, and Simmons 2014; Li et al. 2015; M. Liu, Zhang, and Shen 2014; S. Liu et al. 2015; H. Il Suk, Lee, and Shen 2015; H. Il Suk and Shen 2013a).

Convolutional neural networks (CNNs) are a class of feed-forward artificial neural networks that have absolutely dominated the field of computer vision over the last few years with the success of strikingly superior image classification models based on models including AlexNet (Krizhevsky, Sutskever, and Hinton 2012), ZF Net (Zeiler and Fergus 2014), VGG (Simonyan and Zisserman 2015), GoogleNet (Szegedy et al. 2015), and recently ResNet (He et al. 2016a). Deep CNN models typically stack combinations of convolutional, batch normalization, pooling and rectifier linear (ReLU) operations as a mechanism to reduce the number of connections/parameters in the model while retaining the relevant invariants, and this entire network is typically followed by a fully connected layer that supports inter-node reasoning. The deep residual neural network (ResNet) learning framework as proposed by He et al., 2016a has a similar baseline architecture as the deep CNNs but additionally features parameter-free identity mappings/shortcuts that simplify gradient flow to lower layers during the training phase. Furthermore, each block of layers learns not only from the activations of the preceding block but also from the input to that preceding block. In the original work (He et al. 2016a), these models have been shown to enable ease and simplification of neural network architecture training, thus allowing them to increase network depth and effectively enhance the overall learning performance. These networks radically improve optimization of the “residual” mappings as compared to the collective and unreferenced original mappings (He et al. 2016a) as we will discuss next in more detail in the methods section.

Enhanced performance of the ResNet architecture within the broader imaging community motivated us to explore its diagnostic and prognostic suitability using neuroimaging data in this work. In a systematic approach, we first comprehensively evaluate the diagnostic and prognostic performance of the ResNet architecture implemented in an open-source Pytorch GPU framework (Pytorch Resnet Architecture 2017) on a large dataset (n = 828; see Figure 1 for detailed demographics) featuring cognitively normal (CN), MCI and AD classes. Following this, we focus on the prediction of progression to AD within the MCI class (i.e. predicting which MCI subjects would progress to AD within three years). In this specific analysis, we test the predictive performance of our learning architecture and robustness of the features highlighted by the predictive models, and after that focus on the human brain regions maximally contributing to the prediction of MCI subjects progressing to AD (as suggested by the implemented framework). Finally, we present a qualitative analysis of these results discussing the degree of success (in comparison to previously tested machine learning approaches), limitations and future scope of the evaluated framework to study the diseased brain.

**Figure 1:**
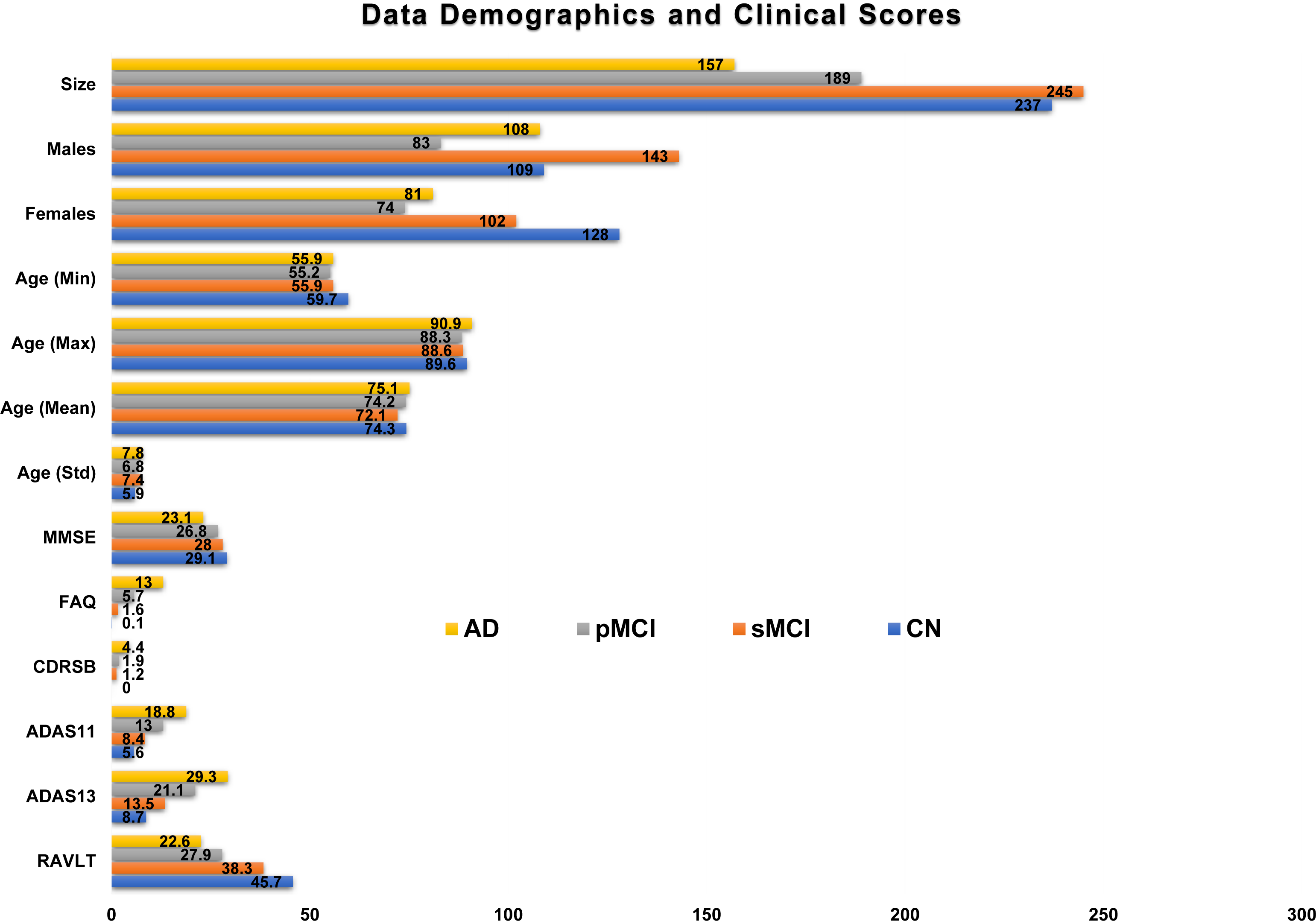
A comparison of data demographics and average clinical scores for the studied classes. This study included all subjects in the ADNI repository that passed the minimum selection criterion (minimum follow-up time, conversion or reversion rules) and pre-processing qualitative analysis. Only the baseline scan for each subject was used for all analyses in this study. Clinical scores for diagnosis: MMSE: Mini-Mental State Exam; FAQ: Functional Activities Questionnaire; CDRSB: Clinical Dementia Rating Sum of Boxes; ADAS: Alzheimer’s Disease Assessment Scale; RAVLT: Rey Auditory Verbal Learning Test.

## Methods

### Structural MRI Data

Data used in the preparation of this article were obtained from the Alzheimer’s Disease Neuroimaging Initiative (ADNI) database (adni.loni.usc.edu). The ADNI was launched in 2003 as a public-private partnership, led by Principal Investigator Michael W. Weiner, MD. The primary goal of ADNI has been to test whether serial magnetic resonance imaging (MRI), positron emission tomography (PET), other biological markers, and clinical and neuropsychological assessment can be combined to measure the progression of mild cognitive impairment (MCI) and early Alzheimer’s disease (AD). For up-to-date information, see www.adni-info.org.

This study worked with all structural MRI scans available in the ADNI 1/2/GO/3 phases (as of November 6, 2017) that passed specific class selection criterion and the image preprocessing pipeline quality check. Healthy aging controls with no conversions in a minimum of 3 years of follow-up from their baseline scans were retained in the cognitively normal (CN) class. Subjects diagnosed as MCI with no conversions/reversions in a minimum of three years of follow-up from their baseline visit were grouped into the stable MCI (sMCI) class, while those converting to AD (multiple conversions excluded) within three years were grouped into the progressive MCI (pMCI) class. Subjects diagnosed as AD at baseline and showing no reversions in a minimum of 2 years of follow-up were retained in the AD class. Only the baseline scan for each subject was used in all analyses. Detailed scanning parameters could be accessed from ADNI data resource webpage (ADNI MRI Protocols). A total number of 830 subjects passed this criterion with further elimination of only two subjects that failed the image preprocessing pipeline quality analysis thus resulting in an overall sample size of 828 subjects for this work. Figure 1 shows the clinical and demographic characteristics of these studied CN, sMCI, pMCI and AD classes.

### Structural Data Pre-processing

Image pre-processing was performed via the statistical parametric mapping 12 (SPM12) toolbox. The structural MRI images were segmented to identify the gray matter brain areas which were spatially normalized and finally smoothed using a 3D Gaussian kernel to 6 mm full width at half maximum (FWHM). The smoothed 3D gray matter images (with a voxel dimension of 160 x 195 x 170) were fed into the deep learning model for diagnostic and prognostic classification. A quality analysis correlation check was conducted with the population mean thresholded image to eliminate outlier (poorly registered) scans. This quality-check discarded only two subjects thus retaining 828 out of the 830 subjects that satisfied the selection criterion which we use for the different diagnostic and prognostic classification tasks in this paper.

### Feature and Class Scores Extraction

A non-linear, deep residual neural network (ResNet) learning framework (He et al. 2016a) was used to extract a series of relatively lower dimensional features from the very high dimensional smoothed 3D images. Similar to the deep CNNs, the deep ResNets are small blocks of multiple convolutional and batch normalization layers followed by a non-linear activation function (typically a rectified linear unit approximated by max(0, x)). While traditional neural networks (NNs) learn to estimate a layer’s or a small stack of layers’ output activation (*y*) as a function (*f*) of the input image or activation (*x*) such that *y* = *f*(*x*), ResNets, on the other hand, feature shortcut identity mappings of input space so as to enable layers to learn incrementally, or in residual representations, with the activations approximated as *y* = *f*(*x*) + *I*(*x*) = *f*(*x*) + *x*, where *I*(∗) is the identity function (He et al. 2016a, 2016b). As such, the latter layers in the ResNets learn not only from the output of the previous layer but also from the input to the preceding residual block, thus gaining extra information at each block in comparison to the traditional NNs. The shortcut connection approach in these networks is similar to that suggested in the “highway networks” (Srivastava, Greff, and Schmidhuber 2015), but differs in being parameter-free (i.e. shortcut connections are identity) as compared to highway networks where shortcut connections are data dependent and parameterized. It has been recently shown (Xie et al. 2017) that the aggregated transformations in this framework allow for substantially stronger representation powers in a homogenous, multi-branched architecture that strikingly requires setting a very small number of hyperparameters. We adapt this model to evaluate the architecture’s performance in pair-wise (binary), mixed-class (domain transfer learning-based binary) and multi-class (4-way) diagnostic classifications as shown in Table 1. While we focus on the progression of the MCI class to the AD class, all other binary classification tasks were undertaken to confirm the appropriateness of learning trends (in terms of classification performance and class separability) in the diagnostic classification of the several disease stages.

**Table 1:** Diagnostic/prognostic classification tasks evaluated through the deep ResNet architecture. Standardized 10-repeat, 5-fold (stratified) cross-validation (CV) framework was employed on each of the mentioned tasks except for the mixed-class task (Task TB below) that varied in that the AD and CN classes were also used for training but only the MCI population was used for testing. Classification task TC corresponds to the multi-class classification task where a four-way classification was performed using the same standardized cross-validation procedure.

| Task | Class 1 | Class 2 | Class 3 | Class 4 | 5-fold stratified CV (10 repeats) |
| --- | --- | --- | --- | --- | --- |
| TA1 | CN | AD | - | - | Standard Binary |
| TA2 | CN | pMCI | - | - | Standard Binary |
| TA3 | sMCI | AD | - | - | Standard Binary |
| TA4 | sMCI | pMCI | - | - | Standard Binary |
| TA5 | CN | sMCI | - | - | Standard Binary |
| TA6 | pMCI | AD | - | - | Standard Binary |
| TB | CN, sMCI | pMCI, AD | - | - | Modified Binary; Split MCI only |
| TC | CN | sMCI | pMCI | AD | Standard 4-way |

In this study, we use a modified form of an open-source Pytorch implementation of this learning framework (Pytorch Resnet Architecture 2017) evaluated for different depths, and reducing the final fully-connected layer to class probabilities to verify classification performance and appropriateness for the studied neuroimaging data. The 3D input data (smoothed gray matter maps) are fed into the deep learning ResNet framework (Figure 2) which has a series of 3D convolutional units (CUs), 3D batch-normalization units (BNUs) and non-linear activation units (Rectifier Linear Units or ReLUs) followed by a max-pooling unit (MPU) from where features are fed to the following residual blocks (RBs). Each RB has two small stacks of layers, also termed building blocks (BBs), with each BB having two CUs, two BNUs and 1 ReLU in the same specific order (CU-BNU-ReLU-CU-BNU). Following the original recommendation (Ioffe and Szegedy 2015) BNUs were adopted following every CU and before any activation functions. The activation at the output of the final residual block adder is fed into an average pooling (AP) unit for dimension reduction and subsequently flattened (from 3D to 1D) to feed a fully connected (FC) layer featuring 512 output nodes. This relatively lower dimensional flattened feature space at the output of the first FC layer (FC1) is fed into a second FC layer (FC2) to estimate the diagnostic class probabilities/scores.

**Figure 2:**
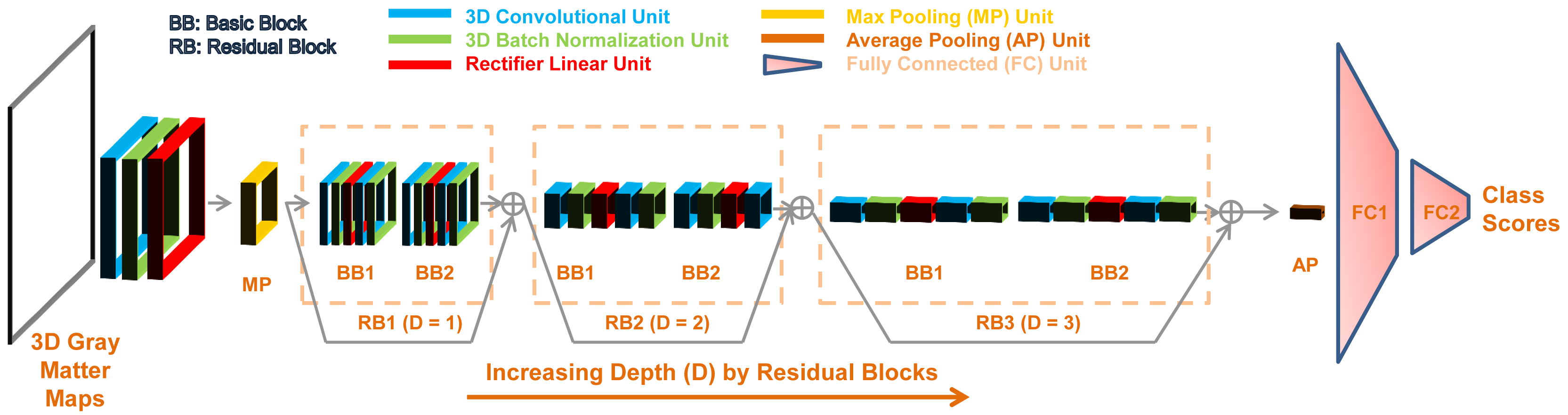
A deep residual neural network learning framework is composed of multiple residual blocks that are small stacks of convolutional and batch normalization layers followed by non-linear activation functions such as rectified linear units. In this study, as suggested by the data (Figure 3), we use a model with three residual layers for evaluating diagnostic classification performance and progression to AD.

Training and testing routines were implemented on an NVIDIA CUDA parallel computing platform (accessing three independent servers each with 4 GeForce GTX 1080 11 GB GPUs) using GPU-accelerated CUDA toolkit/compilation and Pytorch python package tensor libraries. The Adam stochastic optimization algorithm (Kingma and Ba 2015) was preferred for its computational efficiency, relatively low memory requirements, and suitability for problems with large data/parameters size. A batch size of 16, fixed learning rate parameter of 0.001 and L2 weight decay parameter of 0.01 were chosen for the final model selection, and all further classifier performance and feature estimation routines. These settings were based on a preliminary analysis on the CN vs. AD classification task that suggested (1) insignificant effect of batch-size on learner performance, and (2) the validated values of learning rate and L2 weight decay parameter through a grid-search cross-validation analysis. Due to the GPU device memory constraints, we tested only for batch sizes of 2, 4, 8 and 16 and since batch-size did not noticeably affect performance, the maximum batch-size of 16 was chosen to speed up computations (as compared to batch sizes 2, 4 and 8). Subsequently, ResNet’s performance for different model depths (number of residual blocks) was compared to choose the appropriate model depth for consistent comparison across several classification tasks as demonstrated in Table 1.

### Architecture Depth Selection, Regularization and Validation

The ResNet architecture with different depths (*D = 1, 2, 3, 4*; where *D* is the number of residual blocks) was tested for diagnostic classification performance for the CN vs. AD classification task. We retained the architecture depth with the best performance as suggested in this analysis *(D = 3)* for all other classification tasks for consistent comparison. Figure 2 illustrates the modular structure of the selected framework, whereas Figure 3 shows a comparison of the model performances at different depths. As shown in Figure 2, following the MPU, this architecture featured three RBs followed by two FCUs; hence, in all, thirteen convolutional and two fully connected layers were used in this fifteen-layer model. Use of BNUs, default L2 weight decay (regularization) in the Adam Optimizer, repeated stratified k-fold cross-validation for the diagnostic and prognostic classification tasks and early stopping were measures undertaken to prevent any overfitting and reduce classification performance bias. This chosen architecture was then used to extract the features and class probability scores for the different binary/mixed-class/multi-class classification tasks as discussed in the following section.

**Figure 3:**
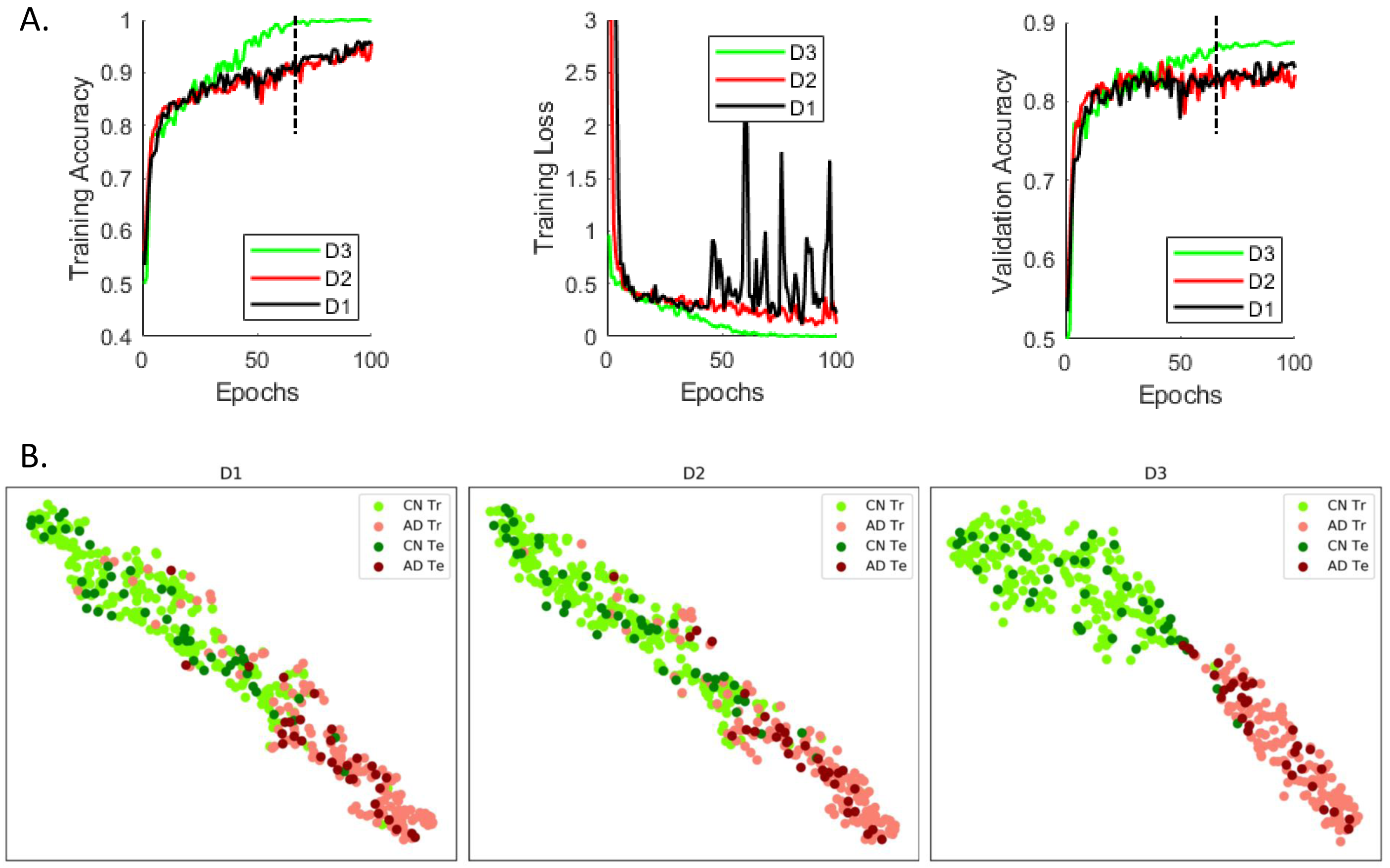
(A) Repeated (n=10) stratified k-fold (k = 5) cross-validation was performed on the pooled cognitively normal (CN) and Alzheimer’s Disease (AD) classes to study the effect of adding depth (i.e. adding further convolutional layers or residual blocks) in the implemented framework. Significant improvement in validation accuracy was reported by a model that used 3 residual blocks (D3: depth = 3) as compared to a model that used 2 residual blocks (D2: depth = 2; p = 1.6996e-07) and a model that used 1 residual block (D1: depth = 1; p = 4.5633e-13). Adding another residual block (i.e. depth = 4) did not result in a significant improvement in performance; hence, we’ve settled on the D3 model and validated it in the several classification/prediction tasks for a consistent comparison. For this specific analysis, all models were run for 100 epochs and used the same training and test datasets in each of the cross-validation folds for consistency in performance comparison. (B) The feature spaces at output of the first fully connected layer in the three surrogate models (for a sample cross-validation fold at the epoch demonstrated by the vertical black line in Figure 3A) were projected onto a two-dimensional space demonstrate additional separation enabled by addition of residual blocks in the ‘D3’ model as compared to the ‘D2’ and ‘D1’ models. The ‘Tr’ abbreviation corresponds to the training samples whereas ‘Te’ corresponds to the samples used to test the learnt model.

### Diagnostic/Prognostic Classification Tasks

Classification performance for the different binary diagnostic and prognostic classification tasks (CN vs. AD, CN vs. pMCI, sMCI vs. AD, sMCI vs. pMCI, CN vs. sMCI, and pMCI vs. AD) for the four studied groups was evaluated (Tasks TA1 through TA6 in Table 1). Additionally, mixed-class inter-MCI (Task TB: CN+sMCI vs. pMCI+AD; training on all CN and AD data plus 80% of sMCI and pMCI data; testing on 20% sMCI and pMCI data) and multi-class (Task TC: 4-way) classification tasks were performed to enhance classification performance and extract additional information than that conveyed by the binary classifiers respectively. Notably, the mixed inter-MCI class classification task was evaluated to explore any other benefits of domain transfer learning (Cheng et al. 2015), i.e. if training the classifier with more data samples (i.e. all CN and AD datasets) resulted in an improvement in the classification performance. While all other classification tasks were conducted to evaluate the framework performance as compared to frameworks used in similar studies in the recent literature, only the mixed/modified inter-MCI classification task was focused on to seek evidence of the most affected brain areas while progressing to AD. All classification tasks were conducted using repeated (n = 10), stratified 5-fold cross-validation procedures on 90% of the subjects to get an estimate of the cross-validated validation accuracy and fold-specific models. The generated models were then tested on the remaining 10% of the subjects to get an estimate of the cross-validated test accuracy. Classification performance metrics including accuracy, sensitivity, specificity, and balanced accuracy were computed and complemented by conducting the receiver operating characteristic (ROC) curve analysis to estimate the area under the curve (AUC) performance metric for the several undertaken classification tasks.

## Results

### Architecture Depth Selection

In a repeated (n=10), stratified 5-fold cross-validation framework, the CN and AD datasets were evaluated for 100 epochs. The stratified cross-validation procedure was performed on the pooled CN and AD classes to study the effect of adding depth to the implemented architecture (i.e. further convolutional layers or residual blocks). This analysis reported significant (p < 0.005) improvement in validation accuracy by a model that used 3 residual blocks (D3: depth = 3) as compared to a model that used 2 residual blocks (D2: depth = 2; *p* = 1.6996e-07) and a model that used 1 residual block (D1: depth = 1; *p* = 4.5633e-13). Adding another residual block (i.e. depth = 4) did not result in significant improvement in performance; hence, we’ve settled on the D3 model and validated it in the several classification/prediction tasks, as will be shown in the forthcoming sub-sections. In this analysis, the models were run for 100 epochs for each depth and used the exact same training and test datasets in each of the cross-validation folds for consistency in performance comparison. A comparison of training error, training loss and validation error for the different depths are shown in Figure 3A. Additionally, the 512-dimensional feature space at the output of the first fully connected layer in the ResNet model was projected onto a two-dimensional space using the t-distributed stochastic neighbor embedding (tSNE) algorithm (der Maaten et al. 2008) to visualize class separation differences with model order. We show projections from a surrogate model (from a sample cross-validation fold) for a sample epoch around which the D3 model clearly exhibits significant differences in validation accuracy (Figure 3B). The projections from other surrogate models (from other cross-validation folds) and other epochs beyond the significant difference showing epoch could be expected to exhibit a similar pattern because of evidence from results in Figure 3A.

### Binary Diagnostic/Prognostic Classification

The performance of the validated (depth = 3) deep learning framework on pair-wise (binary) classification tasks was compared to identify how well the pMCI and AD populations separated from the CN and sMCI populations. These binary classification tasks were conducted using repeated (n = 10), stratified 5-fold cross-validation procedures on 90% of the samples. Model training was conducted with an early stopping with a patience level of 20 epochs (20% of the set maximum number of epochs) to prevent overtraining the validation models. The 10% held out samples were tested on each of the validated fold-specific models. The mean test accuracy for the ResNet architecture for these four classification tasks was found to be statistically significant (*p* < 0.005) over a standard machine learning approach such as the classical support vector machine (SVM) classifier and a standard deep learning approach such as the stacked autoencoder (SAE) as shown in the top panel of Figure 4. The results in bottom panel of Figure 4 reflect a clear trend with the average cross-validation and test metrics for the classification of CN or sMCI classes from pMCI or AD classes distinctly higher than the average metrics for the CN vs. sMCI and pMCI vs. AD classification tasks. Specifically, for the first four classification tasks (CN vs. AD, CN vs. pMCI, sMCI vs. AD, and sMCI vs. pMCI), we report a cross-validated mean validation accuracy of 91.0%, 89.3%, 88.1% and 77.8% respectively and mean test accuracy of 89.3%, 86.5%, 87.5% and 75.1% respectively. The appropriate separability trend across the different classes and genuinely high classification metrics as compared to previous findings in the literature (reviewed recently in (Moradi et al. 2015 and Vieira et al., 2017) in a large heterogeneous sample highlight the suitability of the used deep learning model.

**Figure 4:**
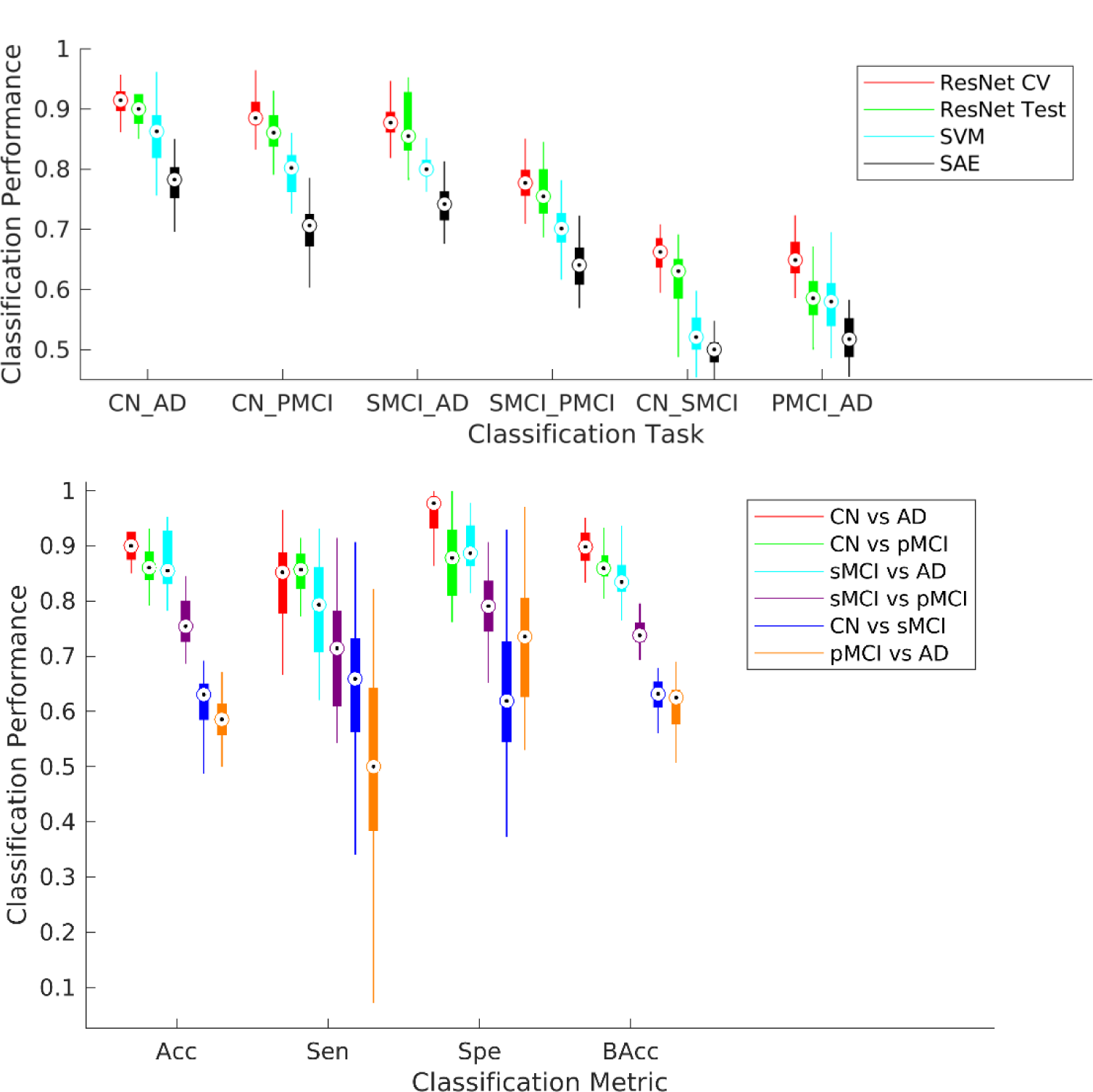
Six possible binary diagnostic and prognostic classification tasks from the four studied classes were considered. A repeated (n = 10), stratified 5-fold cross-validation procedure was conducted for each of these classification tasks. The ResNet framework was trained independently for each classification task for a maximum of 100 epochs but with an early stopping with a patience level of 20 epochs (20% of the set maximum number of epochs) to prevent overtraining the validation models. (Top) The performance of the ResNet framework performed significantly better (*p* < 0.005) than the linear support vector machine (SVM) and stacked auto-encoder (SAE) methods for all binary tasks. (Bottom) Each boxplot shows a spread of the specific reported metric (accuracy, sensitivity, specificity or balanced accuracy) over the 50 cross-validation folds. The first four classification tasks in specific order as in the legend (CN vs. AD, CN vs. pMCI, sMCI vs. AD, and sMCI vs. pMCI) could be considered more clinically relevant and reported a cross-validated mean validation accuracy of 91.0%, 89.3%, 88.1% and 77.8% respectively, and mean test accuracy of 89.3%, 86.5%, 87.5% and 75.1% respectively.

For further introspection into the diagnostic ability of the binary classifiers, we estimated the classification-task-specific receiver operating characteristic (ROC) curves. A comparison of the area under the ROC curve (AUC) metric confirmed a similar trend as suggested in the previous analysis (Figure 4) and as illustrated in Figure 5. We report a cross-validated test AUC of 0.94 for CN vs. AD, 0.90 for CN vs. pMCI, 0.90 for sMCI vs. AD and 0.78 for the sMCI vs. pMCI classification tasks for these tasks respectively. These initial results indicate high suitability of the evaluated framework for our desired objective; further possible improvement in the prediction of progression to AD was explored with the mixed-class prognostic classification analysis as discussed in the next section.

**Figure 5:**
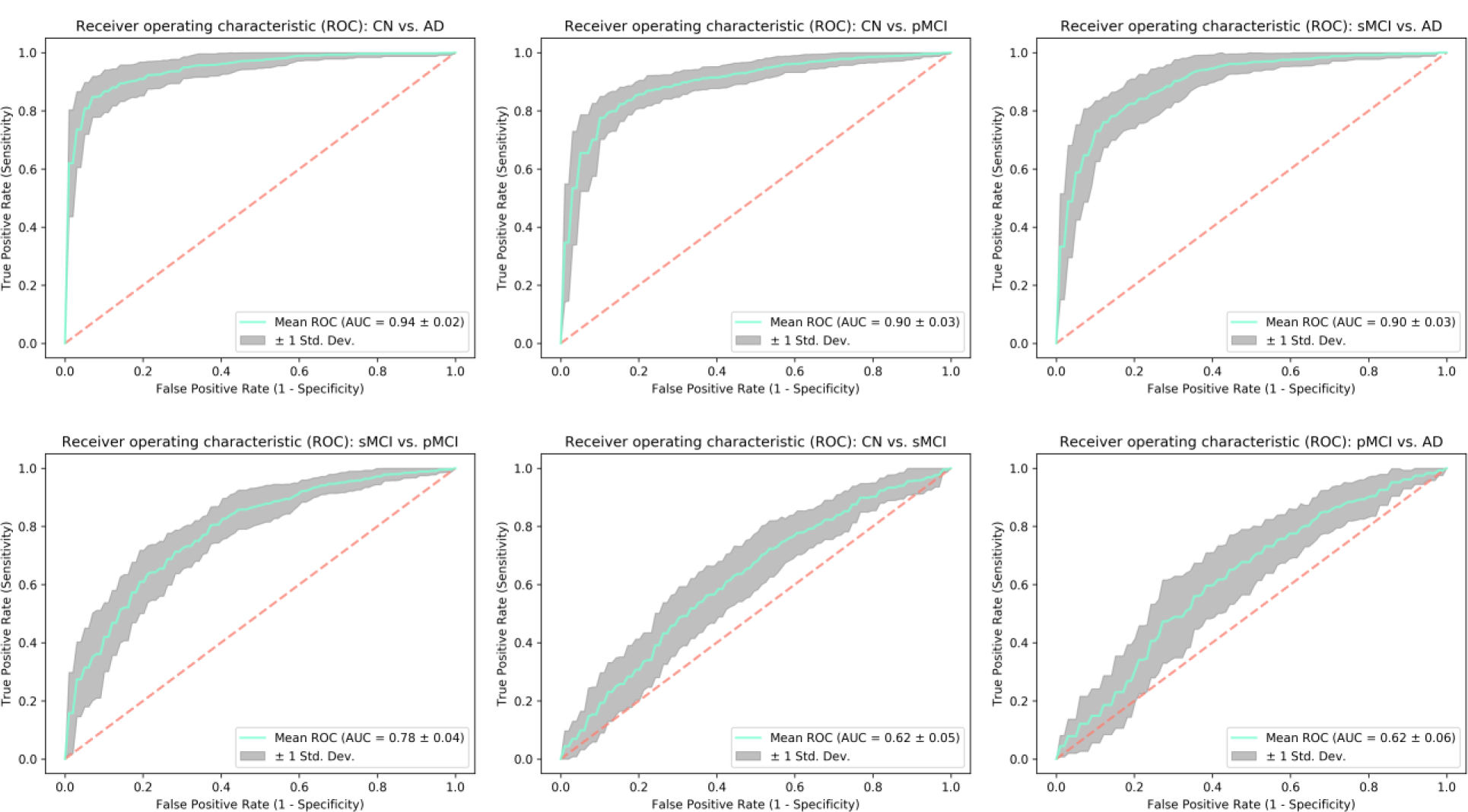
Receiver operating characteristic (ROC) curves were estimated for each of the classification tasks to evaluate the diagnostic ability of the trained ResNet framework further. As expected, the reported area under the curve (AUC) metric follows a similar trend as in Figure 4 thus further adding evidence to the superior performance of the tested architecture for the undertaken analysis.

### Mixed-class Prognostic Classification

The sMCI vs. pMCI classification task could be considered as the most clinically relevant task amongst the several binary classification tasks since identifying MCI subjects who are highly likely to progress to AD is very crucial. Hence, we focus on exploring ways to improve separability between these two classes in this specific analysis. A recent study (Cheng et al. 2015) explored the advantages of domain transfer learning to enhance MCI conversion prediction rates, something similar to what we pursue in the section. In general, training the machine learner with more data is highly likely to improve its classification/prediction performance on unseen data since the learning model assimilates the additional variability provided by the previously unseen datasets and adjusts its weights accordingly for more generalized training (i.e. decrease in generalization error). In a scenario where availability of MCI data is severely limited, we hypothesized that training the learner with all data from the CN and AD classes (or domains) together with some part of the two MCI subtypes (or domains), and then testing with the remaining part of the MCI subtypes (or domains) could enhance classification performance.

For this analysis, we conducted the above discussed modified form of repeated (n=10) stratified 5-fold cross-validation on 100% of the CN and AD class samples, and 90% of the MCI samples (holding the remaining 10% as test samples). We report a significantly improved cross-validated mean test accuracy of 83%, a sensitivity of 76%, a specificity of 87% respectively (Figure 6A), and an AUC of 0.88 respectively (Figure 6B). The results for this modified MCI subtype classification task reflect substantial improvement over the standard binary version of this task (8% in accuracy, 4% in sensitivity, 9% in specificity and 10% in AUC) with the addition of domain transfer learning in the training phase. Finally, the test performance of this modified inter-MCI case was confirmed as a significant improvement (*p* < 0.005) over the standard linear SVM and SAE methods applied on the same training/testing cross-validation folds. In this specific analysis, for estimating the performance of the SVM classifier, the classical univariate feature selection procedure using F-test (ANOVA) was implemented for dimension reduction following which the optimal value of the penalty (cost) parameter in the linear SVM was estimated. For the SAE method, we considered three hidden layers and employed a grid search to select the number of units in the intermediate layers based on the results in H. Il Suk et al., 2015. The boxplots for the accuracies for the different cross-validation folds using the Resnet, SAE and SVM models are shown in Figure 6C. Finally, the cross-validation and test prediction accuracies for the ResNet model on the smoothed gray matter maps are compared to that on the non-smoothed data <u>using the same folds of the stratified, repeated (n = 5) k-fold (k=5) cross-validation</u> (Figure 6D). <u>The performance on the smoothed data was observed to be significantly better (</u>*p* < 0.05) <u>as compared to that evaluated on non-smoothed data. We speculate this resultant improvement due to an increase in the signal to noise ratio caused by smoothing. Notably, smoothing differences were preserved</u>.

**Figure 6:**
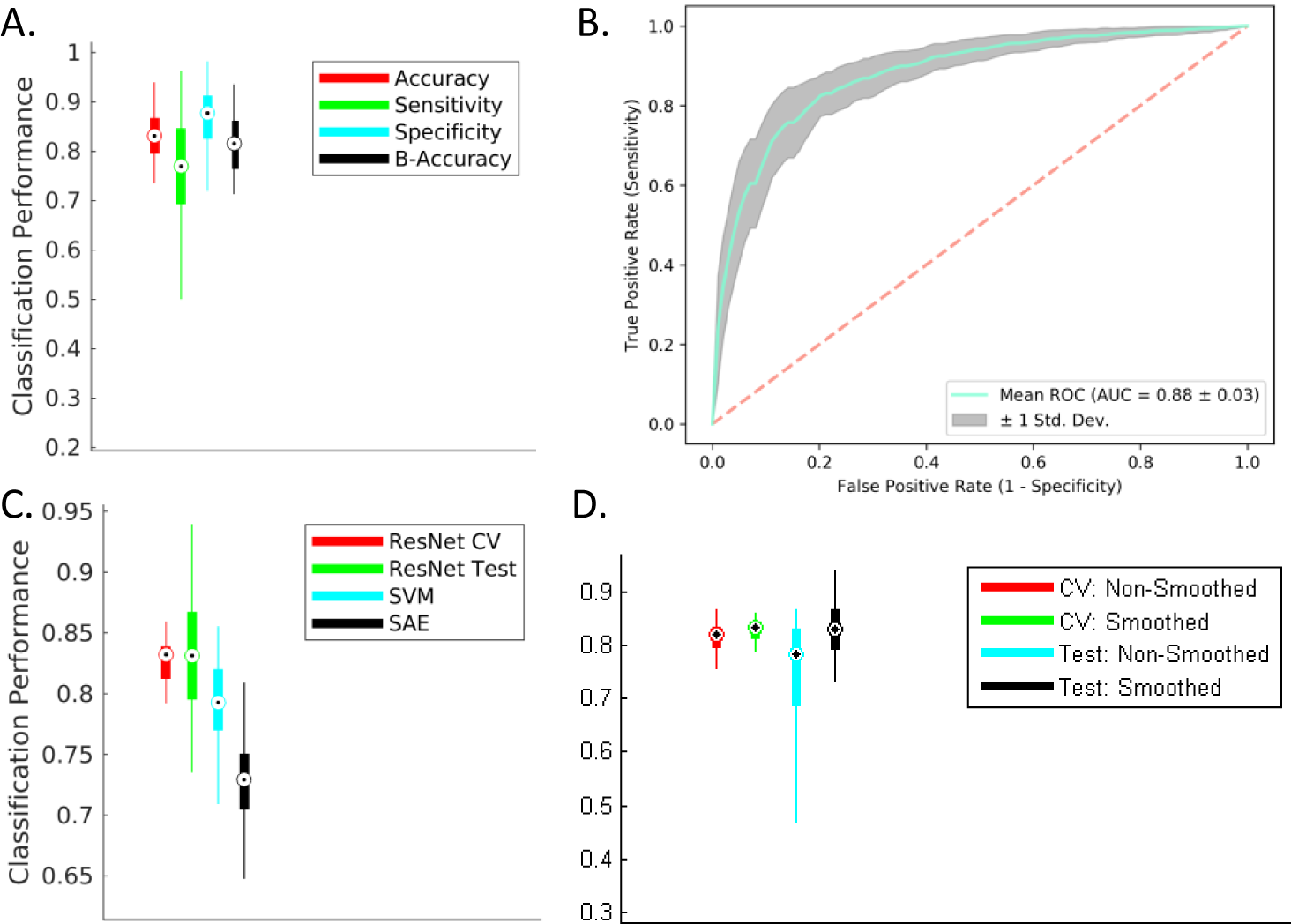
Mixed-Class Prognosis Classification. A modified form of repeated (n = 10), stratified 5-fold cross-validation procedure was conducted to evaluate the separability of the two MCI sub-classes. Hypothesizing an improvement with an increase in amount of training data provided by other classes (analogous to domain transfer learning), the learner was trained with all datasets from the CN and AD classes (or domains) in addition to the cross validation-fold-respective training sMCI/pMCI datasets followed by testing on the cross validation-fold-respective testing sMCI/pMCI datasets. (A) and (B) A significant improvement for all studied classification metrics (6% in accuracy, 7% in sensitivity, 5% in specificity and 7% in AUC) was observed for this mixed-class classification task as compared to the standard inter-MCI class classification task (i.e. sMCI vs. pMCI classification task as shown in Figure 4 and bottom left panel in Figure 5). (C) The mixed-class classification task reported a significant performance improvement (*p* < 0.005) over the classical SVM and SAE methods. (D) The cross-validated validation and test accuracies estimated from the smoothed gray matter maps showed significant improvement (p < 0.05) over the corresponding values estimated from the non-smoothed gray matter maps.

### Comparison with previous literature

In this section, we compare the prediction performance of AD progression in our study (modified inter-MCI task) to previous deep learning work in recent literature (Table 2). Notably, while we directly compare the SAE and SVM methods on the same cross-validation and test folds, a comparison with the results reported from other methods in the reported studies could likely induce a bias due to the indirect nature of comparison (training and testing on different data folds and possibly different conversion periods). Thus, we outstandingly clarify that this section is intended to provide a rich literary review of the most relevant works on AD progression, showcase an indirect comparison to relevant previous works and highlight the potential of the used ResNet in neuroimaging applications such as understanding disease progression. Hence, as such, this comparison doesn’t argue that ResNet architecture is necessarily the most superior of all the frameworks featured in this comparison; rather, our focus is on highlighting the suitability of this architecture in identifying the most discriminative regions and other factors in AD progression.

**Table 2:** Comparison of MCI to AD prediction accuracy using ADNI dataset.

| Study | Sample Size | Conversion Period | Architecture | Cross-validation | Accuracy (%) |
| --- | --- | --- | --- | --- | --- |
| This work | CN = 237<br>sMCI = 245<br>pMCI = 189<br>AD = 157 | 36 months | Residual Neural Network | Repeated (n = 10)<br>Stratified 5-Fold | 83.01 (MRI) |
| H. Il Suk et al., 2015 | CN = 52<br>sMCI = 56<br>pMCI = 43<br>AD = 51 | 18 Months | Stacked Auto-Encoder | Repeated (n = 10)<br>10-Fold | 69.3 (MRI)<br>83.3 (MRI+PET+CSF) |
| H.-I. Suk, Lee, & Shen, 2015 | CN = 52<br>sMCI = 56<br>pMCI = 43<br>AD = 51 | 18 Months | Deep sparse multi-task learning | Repeated (n = 10)<br>10-Fold | 69.8 (MRI)<br>74.2 (MRI+PET) |
|  | CN = 229<br>sMCI = 236<br>pMCI = 167<br>AD = 198 | 18 Months | Deep sparse multi-task learning | Repeated (n = 10)<br>10-Fold | 73.9 (MRI) |
| Li et al., 2015 | CN = 52<br>sMCI = 56<br>pMCI = 43<br>AD = 51 | 18 Months | Multi-layer perceptron | Repeated (n = 10)<br>10-Fold | 57.4 (MRI+PET+CSF) |
| H. Il Suk & Shen, 2013b | CN = 52<br>sMCI = 56<br>pMCI = 43<br>AD = 51 | 18 Months | Stacked Auto-Encoder + Multi-task learning | Repeated (n = 10)<br>10-Fold | 55 (MRI)<br>75.8 (MRI+PET+CSF +SCORES) |
| H.-I. Suk, Lee, Shen, & Initiative, 2017 | CN = 186<br>sMCI = 167<br>pMCI = 226<br>AD = 226 | 18 Months | Multi-Output Linear Regression + Deep Convolution Neural Network (CNN) | Repeated (n = 10)<br>10-Fold | 73.28 (MRI+SCORES) |
|  | -- Same -- |  | Joint Linear and Logistic Regression + Deep CNN | Repeated (n = 10)<br>10-Fold | 74.82 (MRI+SCORES) |
| Shi, Zheng, Li, Zhang, & Ying, 2018 | CN = 52<br>sMCI = 56<br>pMCI = 43<br>AD = 51 | 18 Months | Stacked Deep Polynomial Network | Repeated (n = 5)<br>10-Fold | 78.88 (MRI+PET) |
| H. Il Suk, Lee, & Shen, 2014 | CN = 101<br>sMCI = 128<br>pMCI = 76<br>AD = 93 | Unmentioned | Deep Boltzmann Machine | 10-Fold | 72.42 (MRI)<br>75.92 (MRI+PET) |
| Lu, Popuri, Ding, Balachandar, & Beg, 2018 | CN = 360<br>sMCI = 409<br>pMCI = 217<br>AD = 238 | 0 to 36 Months | Multiscale Deep Neural Network | 10-Fold | 75.44 (MRI)<br>82.93 (MRI+PET) |

To identify previous studies that used deep learning on neuroimaging data to study psychiatric or neurological disorders, we searched PubMed (May 25, 2018) using search terms very similar to a recent review (Vieira, Pinaya, and Mechelli 2017). Specifically, the following search terms were used: (“deep learning” OR “deep architecture” OR “artificial neural network” OR “autoencoder” OR “convolutional neural network” OR “deep belief network”) AND (neurology OR neurological OR psychiatry OR psychiatric OR diagnosis OR prediction OR prognosis OR outcome) AND (neuroimaging OR MRI OR “magnetic resonance imaging” OR “fMRI” OR “functional magnetic resonance imaging” OR PET OR “positron emission tomography”). Following this, we manually screened these articles to identify the relevant subset of studies that applied deep learning to study MCI to AD progression. A comparison of prediction using MRI data only confirms the superior performance of our method as compared to other undertaken approaches. Using just MRI data, the prediction accuracy obtained in our study (83.01%) is numerically 7% greater than the second best performer (using MRI data only) that used a multiscale deep NN in a very recent study (Lu et al. 2018). Considering the use of multiple modalities, only H. Il Suk et al., 2015 (83.3% using MRI, PET and CSF modalities) and Lu et al., 2018 (82.93% using MRI and PET) report slightly higher performance as compared to our study. Interestingly, despite using multiple modalities, the methods used in these two studies report only marginal improvements (0.6% and 0.2% respectively) over our unimodal analysis. Working with multiple modalities generally enhances the prediction performance (variably from 3% to greater than 20% in studies included in Table 2), so it would be reasonable to expect further improvement in prediction performance through our method if complimentary information from an additional modality is leveraged.

Furthermore, Moradi et al., 2015 (see Table 7 in their manuscript) and Korolev et al., 2016 (see Table3 in their manuscript) did extensive comparisons of other (non-deep-learning) studies and showed their respective approaches to result in better precision than other approaches in previous literature. Moradi et al. 2015 studied progression with ADNI data (large sample of 825 subjects) using a regularized logistic regression approach to report classification accuracy of 74% using MRI biomarker only and 82% using their aggregate biomarker that used the patient age and clinical scores as features in addition to the MRI biomarker. Korolev et al. 2016 worked with only ADNI-1 MCI subjects (n = 259) to predict progression to AD from MCI using a probabilistic, kernel-based pattern classification approach to report a prediction accuracy of 79.9% using MRI and clinical (cognitive and functional) scores. Our method predicted more accurately (83.01%) using a large sample of 828 subjects (MRI data alone) than these two multimodal, non-deep-learning studies and all studies reviewed in these two studies.

### Multi-class (4-way) Diagnostic/Prognostic Classification

For the multi-class (4-way) case, the learning framework scored a cross-validated median validation accuracy of 53.8% and test accuracy of 51.41% that is higher than recent studies evaluating such a 4-way classification (as reviewed in Table 2 in Vieira, Pinaya, and Mechelli 2017). However, it must be noted that recent work using traditional pattern recognition approaches has produced superior performances, thus making use of deep learning approaches arguable for such complex classification problems (Sarica, Cerasa, Quattrone, & Calhoun, 2018). As there is ample evidence of the excellent performance of deeper models in complex, generic image processing applications, we note that such a complex 4-way classification problem clearly demands further introspection in the form of extensive hyperparameter validation and choice of the optimization criterion. Nonetheless, the reported accuracy levels are substantially higher than chance (25%). The appropriateness of the data trends learnt in this much harder classification problem was further confirmed by in-depth ROC and feature projection analyses as discussed next. As an extension of binary ROC analysis, for each class, we estimated a single ROC curve by comparing it to all other classes (i.e. one vs all comparison). ROC curves for the multi-class case can also be assessed by micro-averaging which measures true and false positive rates by considering each element of each class as a binary prediction, or by macro-averaging which essentially averages over the several class-specific classification metrics. In this analysis, the AD and CN classes reported a higher AUC of 0.83 and 0.75 respectively, micro-averaged and macro-averaged cases an AUC value of 0.75 and 0.74 respectively, whereas the pMCI and sMCI classes showed lower AUC of 0.71 and 0.68 respectively (Figure 7A).

**Figure 7:**
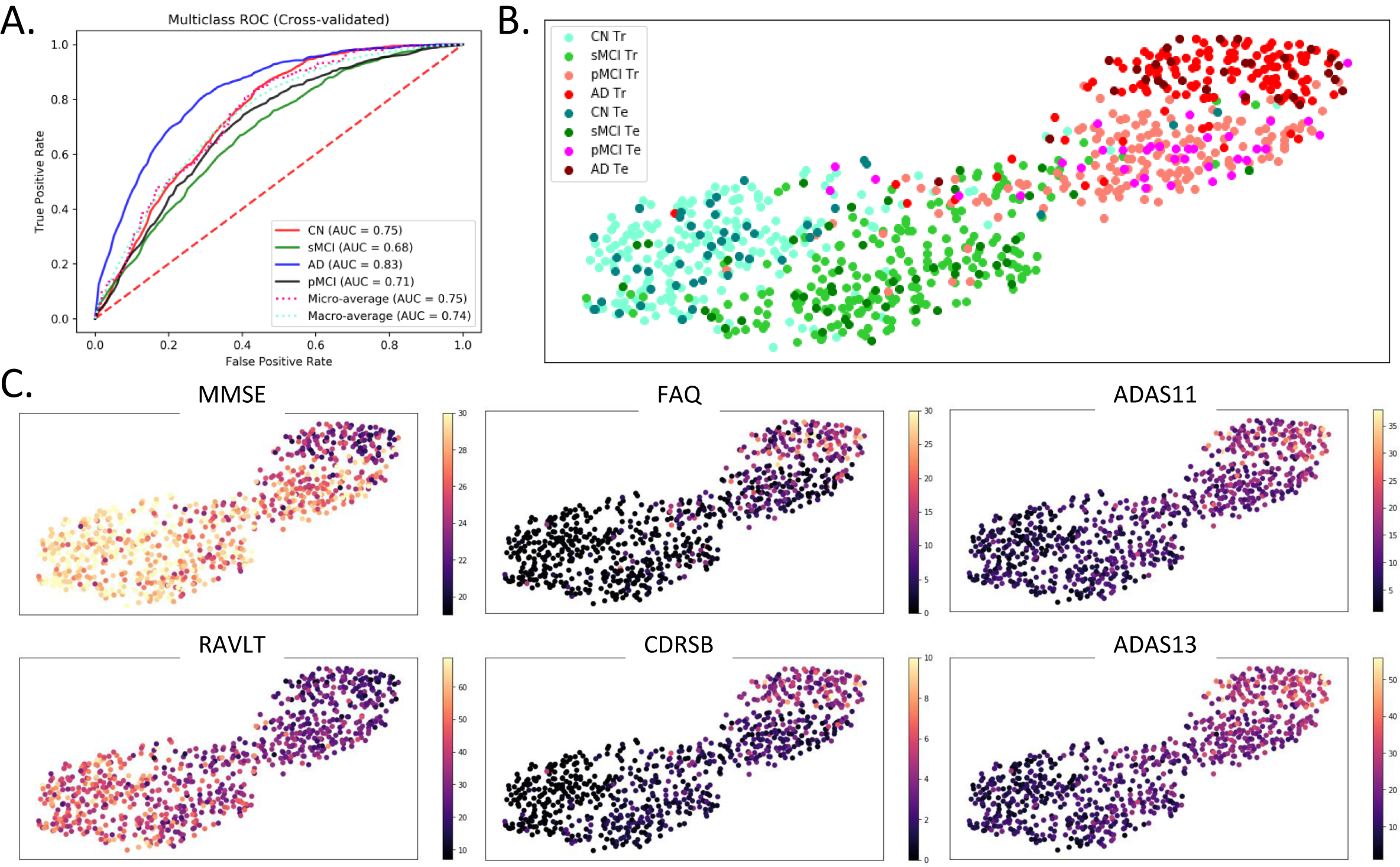
Multi-class ROC and Classification Projection Analysis. (A) For the multi-class classification, ROC analysis for each class was performed by comparing observations from that class to all other classes (i.e. one vs all comparison). Additionally, micro-averaged and macro-averaged ROC estimates were computed to find singular performance metrics for multi-class classification. Higher AUC was reported by the AD and CN classes followed by the micro-averaged and macro-average cases, while both MCI classes reported a lower AUC. (B) and (C) A feature projection analysis was conducted to confirm the appropriateness of the learning directionality in the multi-class classification task. In this analysis, the features at the output of the first fully-connected layer in a sample surrogate multi-class model were projected onto a two-dimensional space using the tSNE algorithm. Barring few outliers, the projections of the observations are appropriately ordered by disease severity in terms of the diagnostic label (panel B) and clinical scores (panel C). In panel B, the ‘Tr’ abbreviation in the figure legend corresponds to the training samples whereas ‘Te’ corresponds to the test samples. In panel C, the following clinical scores were used: - MMSE: Mini-Mental State Exam, FAQ: Functional Activities Questionnaire, CDRSB: Clinical Dementia Rating Sum of Boxes, ADAS: Alzheimer’s Disease Assessment Scale, and RAVLT: Rey Auditory Verbal Learning Test.

In the multi-class feature projection analyses (Figure 7B and 7C), the 512-dimensional features at the output of the first fully-connected layer in the employed framework were projected onto a two-dimensional space using the tSNE algorithm (der Maaten et al. 2008). The tSNE algorithm embeds similar observations as nearby points and non-similar observations as distant points with high probability; so more similar classes could be expected to cluster near each other in the projection space. This projection analysis was performed to confirm the learning directionality of validated models in our multi-class classification case, expecting majority observations for more similar classes being projected/clustered together. Figure 7B demonstrates projections from a sample surrogate model (i.e. model validated for a sample cross-validation fold). Although the classes are not separable in the projection space, yet a clear pattern can be traced easily across the projection spectrum. More specifically, we can observe classes ordered in increasing severity of disease from left to right (i.e. CN, sMCI, pMCI and AD in this specific order) although some outlier observations do exist. The disease severity or the class pattern is further confirmed by coloring the same two-dimensional projections (as in Figure 7B) with the six clinical (cognitive and functional) scores (Figure 7C). The MMSE and RAVLT clinical scores reveal an apparent increase across the spectrum (left through right), whereas the FAQ, CDRSB, ADAS11 and ADAS13 clinical scores (by nature of score characterization) reveal an apparent decrease across the same spectrum.

Interestingly, this projection graph shows the presence of a clear bi-modal structure with most of the CN and sMCI individuals in the first mode and the pMCI and AD individuals in the second mode. So, as a supplementary validation analysis, we focused on the smoothed grey matter maps of the subjects at tSNE extremes and the boundary of the two modes in the tSNE projection. We performed the proposed study by estimating three groups including the homogeneous healthy controls and AD groups at the left and right far ends of the projection spectra (far-CN and far-AD respectively), and a heterogeneous (fused) group in the middle of the projection spectra. Significance testing using t-test statistics was conducted pairwise on preprocessed grey matter maps of these three groups using FDR corrected p-values for a significance level of 0.05. This analysis validated the differences in the input maps for the subjects at the extremes of the tSNE plots (Figure 8). The brain voxels showing significant differences are highlighted by the difference of mean group activations in panels B1, B2 and B3 of this figure. While these differences can be seen clearly for the comparison of the two homogeneous groups in panel B1 of Figure 8, the subjects close to the boundaries separating the modes showed lower significant differences as compared to both homogenous groups (panels B2 and B3 in Figure 8).

**Figure 8:**
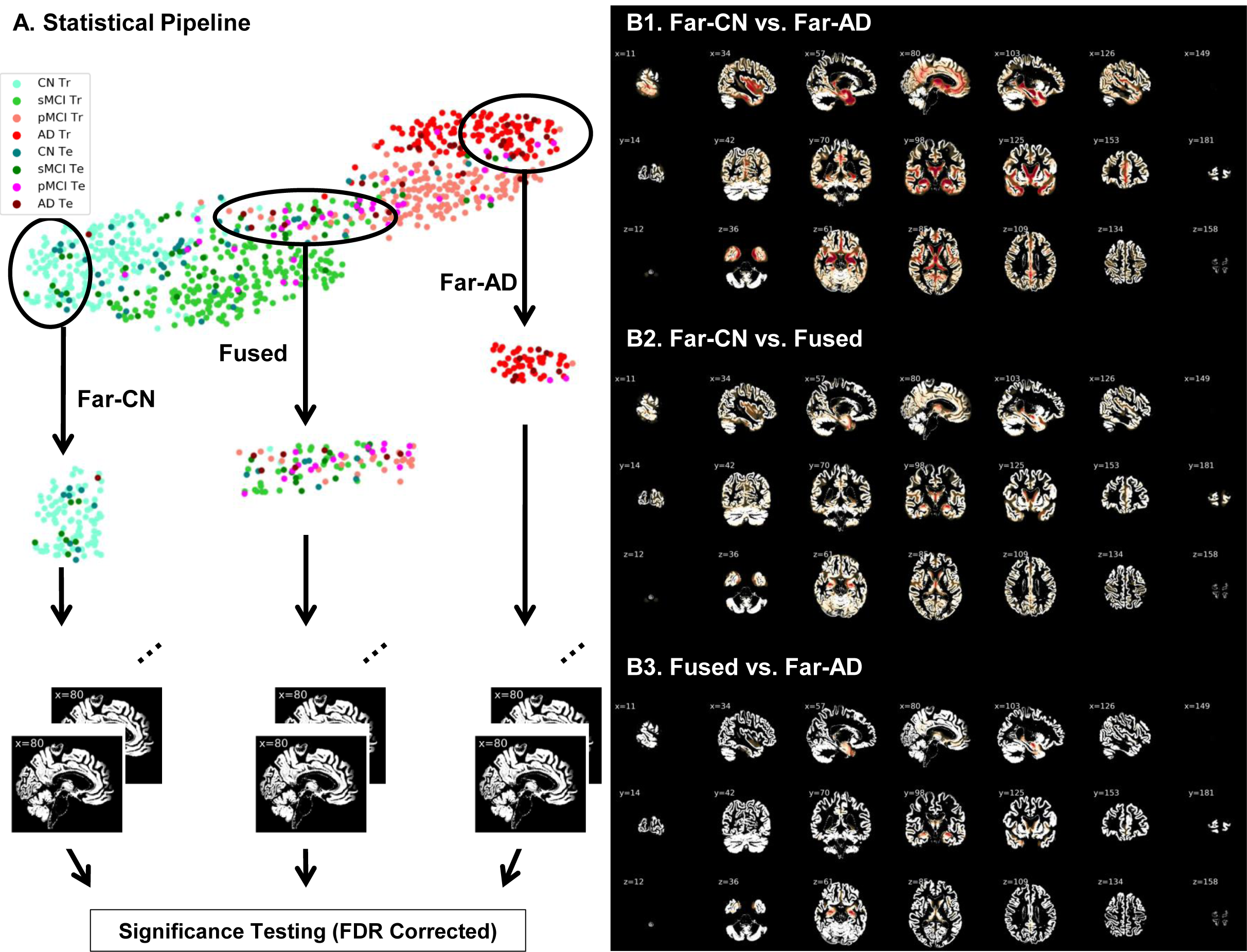
(A) Two-dimensional projections of the 512-dimensional features at the output of the fully connected layer in the ResNet model. Two homogenous groups (far-CN and far-AD) and a heterogeneous group (fused) were sampled and evaluated for significant differences in the input (preprocessed gray matter) space. Voxels showing significant differences post FDR correction (*p* < 0.05) are highlighted in panels B1, B2 and B3. While these differences can be seen clearly for the comparison of the two homogeneous groups in panel B1, the subjects close to the boundaries separating the modes showed lower significant differences as compared to both homogenous groups (panels B2 and B3).

### Localizing Abnormalities: Discriminative Brain Regions

Peak activations of the identified brain regions which are most discriminative of progression of MCI to AD were localized by estimating occlusion sensitivity using the network occlusion approach (Zeiler and Fergus 2014). We pursued this probability-based approach to estimate and quantify the relevance of the different brain regions in the classification decisions, although few other popular approaches (Nair, Precup, Arnold, & Arbel, 2018; Zintgraf, Cohen, Adel, & Welling, 2017) could be adapted too. In this approach, brain networks in correspondence with the automated anatomical labelling (AAL) brain atlas were occluded one at a time, class probabilities re-evaluated, and the relevance of each brain region was estimated proportional to the decrease in target class probabilities when that specific region was occluded. The most discriminative brain networks highlighted through this approach are illustrated in Figure 9A. Peak activations (i.e. highest relevance weights) were observed in the hippocampus, parahippocampal gyrus, temporal superior, middle and inferior gyrus, fusiform gyrus, occipital superior, middle and inferior gyrus including calcarine and cuneus, lingual gyrus, frontal middle and inferior gyrus regions, precuneus, and cerebellum 6, crux 1 and 2 regions. Besides, the amygdala, putamen, thalamus, caudate and frontal superior regions showed moderate relevance.

**Figure 9:**
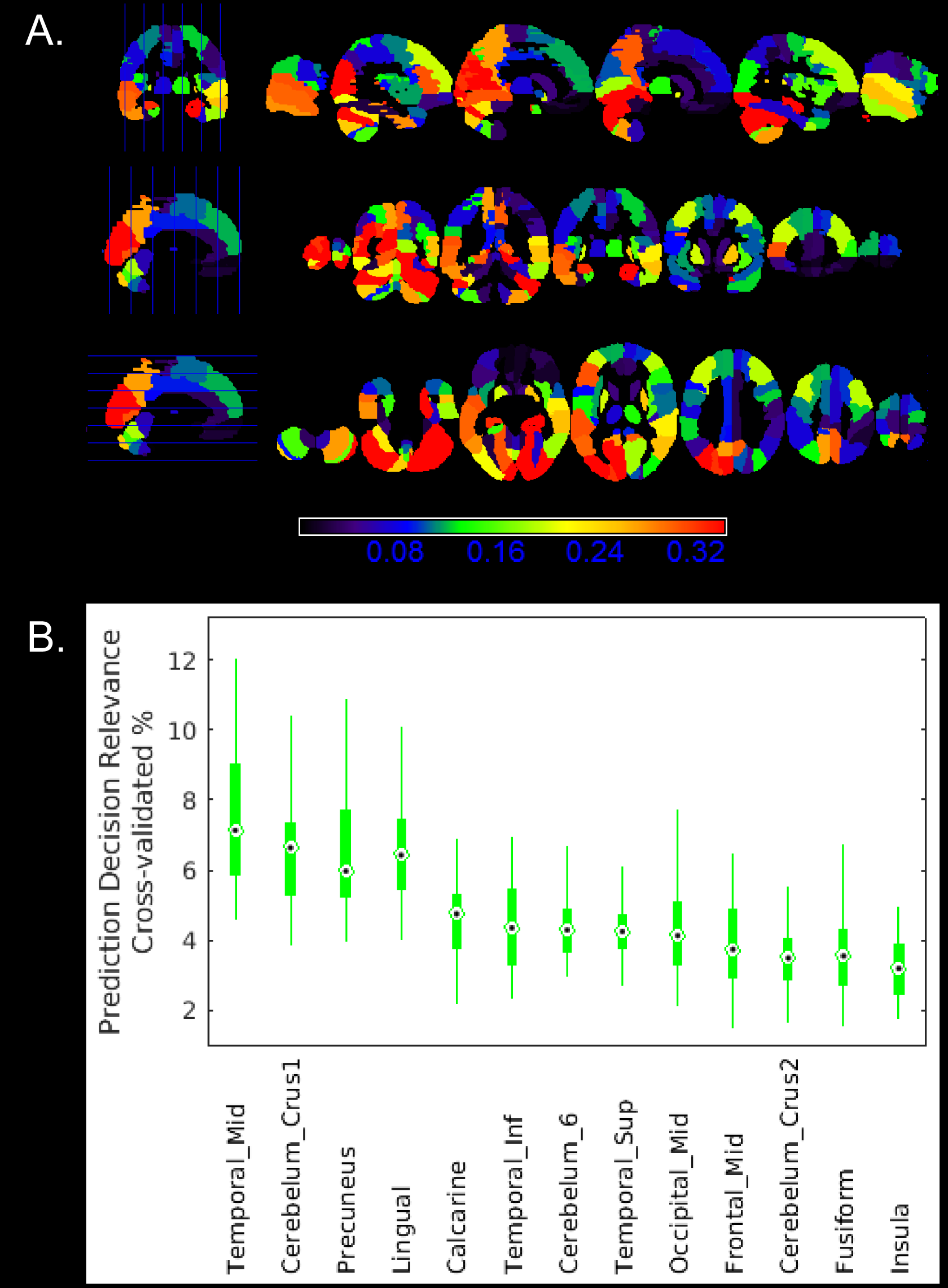
(A) Sagittal, coronal and axial slices of whole brain relevance maps as highlighted by the network occlusion approach in correspondence to the AAL brain atlas networks. (B) Quantitative (cross-validated) assessment of the relevance of the brain regions in classification/prediction decisions to study AD progression. This latter assessment factored in the brain network areas for relevance estimation.

Furthermore, we quantify network relevance estimates by factoring in the network areas (in addition to the assessed change in probabilities). Figure 9B shows the cross-validated percentage contribution of each of the highly relevant networks to the prediction decision making. The illustrated brain regions are the thirteen (out of a total number of 116 AAL brain regions) that consistently emerged in the top 20 most relevant regions in all cross-validation folds. Specifically, highest relevance weights through this latter approach were observed in temporal middle gyrus, cerebellum crus1, precuneus, lingual gyrus and calcarine brain regions, followed by high relevance weights in the temporal inferior gyrus, cerebellum 6, temporal superior gyrus, occipital middle gyrus, frontal middle gyrus, cerebellum 2, fusiform gyrus and insula regions as shown in Figure 9B.

## Discussion

In this work, we extensively test the ability of the ResNets to learn abstract neuroanatomical alterations in structural MRI data. For each of the binary as well as the mixed class (modified inter-MCI) classification tasks, the ResNet architecture performed superior to the SVM and SAE methods. The primary progression analysis of our work is the mixed-class inter-MCI classification task where we used principles of domain transfer learning (additionally training with data from other domains). This analysis bears high clinical relevance. Importantly, on the MRI data alone we achieved a test classification accuracy of 83.01% which is a significant improvement over state of the art with either MRI based (75.44% as reported in Lu et al., 2018) and very close to state of the art performance with multimodal results (83.3% as reported in H. Il Suk et al., 2015). The accuracy in this modified inter-MCI class classification task is significantly higher than that in the standard inter-MCI case which suggests the performance improvement was also enabled by additional training information acquired from other (AD and CN) domains. Notably, the reported performance metrics were obtained from a large dataset (n = 828), a rigorous cross-validation procedure featuring ten repeats and a sufficiently large (20%) validation size and test (10%) size.

Furthermore, the learning directionality and trends were verified in the multiclass case by projecting the features at the output of the first fully-connected layer onto a two-dimensional surface. The projection/clustering class sequence in Figure 7B and 7C support the appropriateness of the extracted features and their association with the clinical scores, thus confirming the high learning capacity and potential of this deep architecture. These results manifest that the ResNets can be considered well-suited to neuroimaging data and future studies to uncover the further potential of such or similar architectures must be undertaken. Next, we discuss the discriminative brain regions suggested by the ResNet in context to previous findings in the literature.

### Discriminative brain regions

AD is characterized by severe trouble in performing familiar tasks, solving problems, planning, reasoning, judgement and thinking, and generally features increased confusion and discomfort in speech, vision, reading, focusing, and spatial or temporal perception. Struggling with these symptoms, the person undergoes mood and personality changes and increasingly loses interest in favorite activities and social life. A sizable amount of previous work has related the above mentioned cognitive, behavioral and emotional phenomenon to specific structural changes in the brain, which we discuss next in context to the discriminative brain regions identified by the ResNet framework.

The hippocampus and amygdala subcortical regions in the medial temporal lobe have been consistently reported as most prominent discriminative regions in early AD. Hippocampus is strongly related to memory formation and recall, and recent evidence suggests more pronounced hippocampal atrophy in the progressive MCI class (Braak and Braak 1991b; Burton et al. 2009; Costafreda et al. 2011; Devanand et al. 2007; Kantarci et al. 2009; Risacher et al. 2009; Visser et al. 2002; Walhovd et al. 2010). Similarly, structural changes in the amygdala, a brain region mainly responsible for emotional experiences and expressions, have been related to personality changes, for example, increased irritability and anxiety, in AD (Poulin et al. 2011; Unger et al. 1991; Whitwell et al. 2008). Other relatively highly activated subcortical regions included para-hippocampal gyrus, thalamus and putamen. While the primary function of the thalamus is to relay motor and sensory signals to the cerebral cortex and regulate consciousness and sleep, the dorsal striatum is believed to contribute directly to decision-making subjective to desired goals. Observed aberrations in the putamen and thalamus regions are typical of subjects with AD (Aggleton et al. 2016; Braak and Braak 1991a; Cho et al. 2014; Jiji et al. 2013; De Jong et al. 2008). Impairments in the thalamus in AD have been associated with deteriorating consciousness, bodily movement and coordination, attentional, and motivation levels and impairments in the dorsal striatum related to very slow or absent decision-making abilities.

Apart from the above widely studied and highly discriminative medial temporal lobe, we also report peak activations in the inferior and superior temporal gyruses and the fusiform gyrus. These regions have been known to be associated with pattern (e.g. face, body, object, word, color, etc.) recognition and reported to be affected by AD in a few previous studies (Chan et al. 2001; Galton et al. 2011). In the frontal lobe, peak activations were observed in the middle and inferior frontal gyrus. These regions are also associated with decision making and problem-solving, reportedly highly damaged in AD (Johnson et al. 2005; Sluimer et al. 2009; Whitwell et al. 2008) and are believed to lead to higher lethargy levels, bizarre/inappropriate behavior and situations of being stuck in a specific condition (repeating same things over and over again).

Besides the above discussed frontotemporal networks, AD is characterized by a decline in critical parietal networks such as precuneus (Apostolova and Thompson 2008; Bailly et al. 2015; Fennema-Notestine et al. 2009; Scahill et al. 2002; Walhovd et al. 2010; Whitwell et al. 2008). Cerebellum, a critical brain region in several motor, cognitive and behavioral functions, is also more recently being increasingly suggested as a direct contributor to cognitive and neuropsychiatric deficits in AD (Guo et al. 2016; Jacobs et al. 2017; Schmahmann 2016). Deteriorating cerebellum health resulting in several symptoms such as lack of balance and coordination, tremors, slurred speech and abnormal eye movements in the elderly. Finally, damages to the occipital lobe are associated with increased misinterpretations of the surrounding environment (e.g. hallucinations, illusions, misidentification, misperceptions, etc.) and occipital regions comprising the calcarine, cuneus and lingual gyrus regions have indeed been reported to be compromised in progression to AD.

The above-discussed findings add further evidence that the localized abnormal patterns in the brain structure could play a significant role in the prediction of early AD biomarkers and are of potential clinical application. A few of the discriminative regions that we report are rarely used as prognostic biomarkers to study the conversion of MCI to AD; our work and the cited literature in this discussion provide compelling evidence of including these new biomarkers to allow for a complete characterization of the structural changes in AD progression.

### Limitations and future scope

Here we note some inherent limitations of our work that could be addressed in the future depending on algorithmic computational tractability and availability of data resources and data processing algorithms. As with other neuroimaging studies, the foremost limitation is a limited training data size. In generic image processing applications, this limitation is often addressed with data augmentation procedures by using simple rotation, translation, scaling and other data transformations (also see Castro et al., 2015 and Ulloa et al., 2015 for more elaborate data augmentation examples with structural MRI). We expect even further increases in performance by employing such techniques in future work, a fact that broadens perspectives for our models that are already performing at or above state of the art.

Interestingly, a recent study (Casanova, Hsu, and Espeland 2012) demonstrated an increase in classification performance with an increase in sample size using ADNI structural MRI data. Similarly, in our work as well, we saw a substantial increase in performance with more training data being fed to the ResNet framework in the modified inter-MCI class classification task as compared to the standard inter-MCI class classification task. This makes a strong case to test the use of multiple datasets to extract features in a pooled or separate fashion and then use the pooled or separate information to train the machine learning framework. With increasing data availability and standardization in data preprocessing and pooling procedures, further substantial improvement in diagnostic and prognostic classification performance could be expected in future multi-study deep learning research efforts.

Due to the computationally expensive nature of training deep CNNs, few limitations regarding computational tractability within realistic study time remain. This tends to restrict extensive fine-tuning of each involved hyperparameter through random or grid search analysis on multiple hyperparameters and additionally backing up statistical trends using methods such as Monte-Carlo. As such, the most critical hyperparameters must be prioritized and optimized to estimate general performance trends of the algorithm within the realistic study period. For this specific work, we optimized the initial learning rate and L2 weight decay parameter on a sample cross-validation fold using extensive grid analysis and retained the values for other dataset partitions. Although the same hyperparameters would likely achieve close to actual performance on different data folds, yet this fine-tuning could have a small effect on the performance of the respective surrogate models (e.g. reported performance metrics could be slightly lower than the original) but also on that of the final predictive model. It must be noted that this limitation is for performance quantification only; it is least likely to affect the qualitative analysis (e.g. localizing discriminative brain regions) by a significant margin.

Choosing a stopping criterion for learning a classifier typically involves a tradeoff between generalization error and learning time. While this study approximated the stopping criterion with information across all cross-validation folds, further detailed introspection using relatively unestablished but promising variants of early-stopping criterion could be explored in future investigations (Prechelt 1998). Similarly, the effect of algorithmic variations in bottleneck residual block structures (size and depth), training time, and loss optimization procedures could be understood in future studies to enhance the prediction performance further.

Several other approaches for enhancing predictive performance of AD progression could be explored in future work. Diagnosis for the subjects is currently established through clinical scores, but diagnosis-specific neuroanatomical or neurofunctional abnormalities might not show in each subject in each class due to the heterogeneous nature of age-related dementia. In such a scenario, it could be interesting to constrain this heterogeneity by training the machine learning model on the most homogeneous samples (i.e. samples most representative of the given class) and then evaluate the change in the performance of the diagnostic/prognostic classification or change in the feature space of interest. Another approach could be to fuse the low-dimensional clinical scores used to make the clinical diagnosis with the MRI features space to further enrich the feature learning process. This approach has reportedly resulted in enhanced performance in few studies as also suggested in Table 2. Various widely used low dimensional features chosen by experts (e.g. volumetric MRI features or similar features from other modalities) could even further enhance diagnostic and prognostic performance.

Recent literature reflects ample evidence of advantages of multimodal studies in understanding brain structure and function and decodes brain complexities (Abrol, Rashid, et al. 2017; Calhoun and Adali 2009; Calhoun and Sui 2016). Indeed, few previous multimodal studies have reported prediction performance improvements due to training the same machine learning framework with multiple modalities as compared to a single studied modality for studying AD/MCI (Lorenzi et al. 2016; Toledo et al. 2013; Zhang et al. 2011) as is also evident from Table 2. Due to this evidence from other explored machine learning neuroimaging studies, performance improvement is highly likely if features for multiple modalities are extracted through the ResNet framework and further fused using a data fusion algorithm to generate a collective feature space for predicting chances of progression to AD.

Interestingly, the fusion of features from multiple structural (MRI, PET and CSF) modalities (structure-structure fusion) has been much more frequently explored than the fusion of feature space from one or more of these structural modalities to feature space from a functional neuroimaging modality (structure-function fusion) such as fMRI. One of the reasons for the relatively less explored structure-function fusion in AD/MCI literature could be the significantly smaller number of fMRI datasets as compared to data from the structural modalities. Nonetheless, structure-function fusion could be highly engaging, and several robust fMRI features such as amplitude of low-frequency fluctuation (ALFF) maps, or static/time-varying functional connectivity (FC) maps exist. Of specific interest in fMRI is the time-varying FC feature space that have recently been shown to be replicable (Abrol et al. 2016), statistically significant and robust against variation in data grouping, decomposition and analysis methods (Abrol, Damaraju, et al. 2017), and also more discriminative of diseased brain conditions (Rashid et al. 2016) than static FC. As such, future works featuring such promising deep learning models could seek performance gains not only from structure-structure fusion coupled with information in cognitive/functional scores but with structure-function fusion as well.

## Conclusion

This work shows that the ResNet architecture showed performance numerically comparable to state of the art in predicting progression to AD using MRI data alone, and within 1% of the state of art performance considering multimodal studies as well. This clearly reflects the high potential of this deep architecture for studying progression to AD and neuroimaging data in a broader sense. The prognostic classification performance was exceptional despite several limitations as outlined in the discussion section and addressing these limitations in future work could highly likely result in further improvement in the performance of this relatively newer machine learning framework. The most discriminative brain regions as highlighted by the ResNet framework confer with previous findings in AD/MCI literature to a high degree, and brain regions for which there is insufficient evidence must be investigated further to enhance the set of potential AD biomarkers. The ResNet architecture could be explored in future for learning from multiple modalities for examining any possible improvements in diagnostic and prognostic classification and identification of more specific multi-modal biomarkers for AD or other brain conditions. We conclude that our results further strengthen the expectations and a high likelihood of discovery of modifiable risk factors for understanding biomarkers of progression to AD early, primarily using advanced neuroimaging data processing methods such as the one explored in this work.

## Data Availability Statement

The data (pipeline and generated deep learning models) that support the findings of this study are available from the corresponding author (Dr. Anees Abrol) upon reasonable request.

## ACKNOWLEDGEMENT

This work was supported by NIH grant numbers 2R01EB005846, P20GM103472, and R01REB020407 as well as NSF grant 1539067 to Dr. Vince D. Calhoun, as well as National Natural Science Foundation of China grant 61703253, Natural Science Foundation of Shanxi Province in China grant 2016021077 to Dr. Yuhui Du and NSF grant IIS-1318759 to Dr. Sergey Plis.

Data collection and sharing for this project was funded by the Alzheimer’s Disease Neuroimaging Initiative (ADNI) (National Institutes of Health Grant U01 AG024904) and DOD ADNI (Department of Defense award number W81XWH-12-2-0012). ADNI is funded by the National Institute on Aging, the National Institute of Biomedical Imaging and Bioengineering, and through generous contributions from the following: AbbVie, Alzheimer’s Association; Alzheimer’s Drug Discovery Foundation; Araclon Biotech; BioClinica, Inc.; Biogen; Bristol-Myers Squibb Company; CereSpir, Inc.; Cogstate; Eisai Inc.; Elan Pharmaceuticals, Inc.; Eli Lilly and Company; EuroImmun; F. Hoffmann-La Roche Ltd and its affiliated company Genentech, Inc.; Fujirebio; GE Healthcare; IXICO Ltd.; Janssen Alzheimer Immunotherapy Research & Development, LLC.; Johnson & Johnson Pharmaceutical Research & Development LLC.; Lumosity; Lundbeck; Merck & Co., Inc.; Meso Scale Diagnostics, LLC.; NeuroRx Research; Neurotrack Technologies; Novartis Pharmaceuticals Corporation; Pfizer Inc.; Piramal Imaging; Servier; Takeda Pharmaceutical Company; and Transition Therapeutics. The Canadian Institutes of Health Research is providing funds to support ADNI clinical sites in Canada. Private sector contributions are facilitated by the Foundation for the National Institutes of Health (www.fnih.org). The grantee organization is the Northern California Institute for Research and Education, and the study is coordinated by the Alzheimer’s Therapeutic Research Institute at the University of Southern California. ADNI data are disseminated by the Laboratory for Neuro Imaging at the University of Southern California.

